# Emergent spatial structure in the gut microbiota is driven by bacterial growth and gut contractions

**DOI:** 10.1101/2025.05.22.655445

**Authors:** Giorgia Greter, Sebastian Hummel, Daria Künzli, Naomi Dünki, Niina Ruoho, Patricia Burkhardt, Suwannee Ganguillet, Milad Radiom, Claudia Moresi, Leanid Laganenka, Wolf-Dietrich Hardt, Steffen Geisel, Sebastian Jordi, Benjamin Misselwitz, Bahtiyar Yilmaz, Jonasz Slomka, Eleonora Secchi, Roman Stocker, Emma Slack, Markus Arnoldini

## Abstract

Spatial structure can determine function and evolution of bacterial communities. The gut microbiota is known to be spatially structured longitudinally along the many meters of the gastrointestinal tract, but micro-scale structure in the gut lumen has not been extensively explored. In samples from mice and humans, we show that upper large-intestinal content behaves as a non-Newtonian fluid that changes its viscoelastic properties under the force of gut contractions. This phenomenon is sufficient to explain micro-scale bacterial clustering in the murine cecum, resulting from growth within the gel-like structure of cecum content, and periodic disruption due to peristalsis-driven shear-thinning and clearance. Shear-thinning can also explain the surprising observation that fed beads enter the tip of the mouse cecum by flow along the epithelial cell layer before being mixed into the cecum content. Our study shows mechanistically how spatial structure in the gut emerges through the interplay of microbial and host physiology and highlights the possibility of host control over gut microbiota distribution via gut contractions.

**One sentence summary:** We show how spatial structure emerges in the gut microbiota through bacterial growth in the matrix of gut content.

## Introduction

The gastrointestinal microbiota plays important roles in maintaining host health and metabolic homeostasis (*1, 2*). Along the gastrointestinal tract, microbiota density and composition vary due to changing environmental conditions, such as pH, flow rates and oxygen levels, that microbes encounter along its length (*3–6*). In the dense bacterial community in the large intestine, bacteria are not well mixed, even at length scales where such global parameters are constant (*7–11*). Given the short ranges at which positive (*12*) and negative (*13*) interactions between bacteria can take place, the direct neighborhood of bacteria is important in determining how they live. Understanding the extent and origins of spatial structure in the gut microbiota is thus crucial for mechanistically understanding ecological and evolutionary processes in this important microbial community.

There are different possible mechanisms for how spatial structure in the large intestine could emerge. Anatomical features, such as colonic crypts, the mucus layer in the distal colon, and oxygen gradients close to the epithelium can create spatially distinct habitats (*10, 14–17*), but the bulk of bacterial growth happens in the gut lumen of the proximal large intestine without such pre-defined structural features (*6, 9, 18*). Nevertheless, micro-scale spatial structure can emerge in the gut lumen, and possible mechanisms include small-scale differences in physical properties (e.g. density, viscosity, adherence to food particles) of luminal content (*19–21*), cross-linking of bacteria by antibodies (*22*), and flagella-driven chemotactic movement of bacteria (*23, 24*). If the environment prevents dispersal and nutrients are available, bacterial cell division can generate patches of clonal cells (*25*). The size of such microcolonies will depend on the niche size, as bacterial growth depends on the ecological space and resources available in a niche, as well as the stability of that niche (*26–28*).

All the processes that can lead to spatial structure in the gut lumen take place in, and interact with, the complex matrix of gut content. Gut contractions, which play an important role in maintaining microbial abundance and composition (*6, 29, 30*), exert forces on the gut lumen when propelling and mixing luminal content. Intestinal content is shear thinning (i.e., it displays decreasing viscosity with increasing shear rate), and changes its properties from more solid-like to more liquid-like when a contractile force is applied (i.e., it undergoes a yielding transition) (*31, 32*). This may render bacterial swimming, nutrient distribution, and more generally the possibility of forming spatial structure in the lumen dependent on gut contractions: particulates and cells can move more easily when gut content behaves liquid-like, while they are kept in place when it behaves solid-like.

Here, we quantify the influence of different possible mechanisms governing bacterial spatial dynamics in the mammalian gut. While chemotaxis and antibody binding play detectable roles, we conclude that bacterial growth is a major factor in determining bacterial clustering in the gut. In the gel-like matrix of gut content that restricts movement, bacterial cell division results in microcolony formation, and the extent of microcolony formation is limited by gut contractions (*7–11*). By experimentally modulating flow and mixing in the gut lumen of mice, and by quantifying rheological parameters of gut content from mice and humans, we reveal how the host can change the biophysical properties of the gut lumen via gut contractions, potentially giving it control over the extent of microcolony formation and thus ecological interactions in the gut microbiota.

## Results

### Organ-scale spatiotemporal dynamics of mixing in the murine cecum

To experimentally assess the strength and dynamics of mixing in the mouse cecum, the part of the large intestine with most bacterial activity, we have analyzed the distribution of bacteria in two parts of the organ: at the cecum tip (the part of the cecum furthest away from the ileum and colon) and the cecum base (the part of the cecum close to where ileum and colon connect to it) (**Figure 1A**). To allow full visualization of the distribution of different microbiota members, we used a gnotobiotic mouse model containing three bacterial species (three-member microbiota, 3MM (*33*)), *Bacteroides thetaiotaomicron* (*B. theta*), *Eubacterium rectale* (*E. rectale*), and *Escherichia coli* (*E. coli*) (**Figure 1B**). Bacteria in fixed cryosections of cecum tissue and content were visualized using fluorescent *in-situ* hybridization (FISH) and confocal microscopy, and center coordinates for every bacterium were determined using a custom image analysis method (**Figure S1**, Methods).

**Figure 1:**
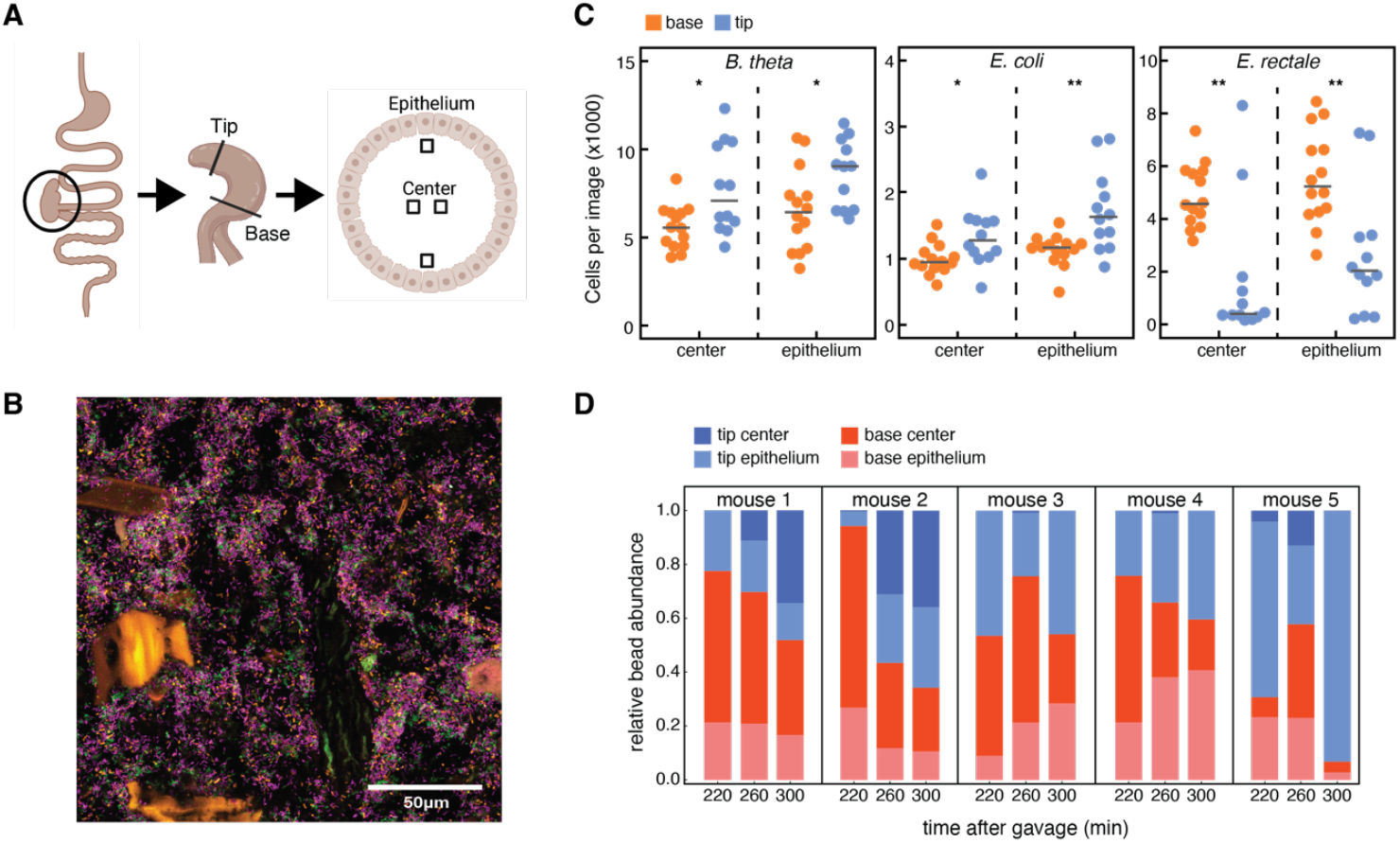
Bacteria are not homogenously mixed throughout the murine cecum. (**A**) Schematic illustrating the experimental procedure for capturing images. The entire cecum is fixed and frozen, and cryosections are produced for tip and base. Four images were taken for each section: two at the center and two at the epithelium. (**B**) Representative microscopy image of the 3MM gnotobiotic model taken at the center base. Green cells are *B. theta*, orange cells are *E. coli*, purple cells are *E. rectale*. Autofluorescent large food particles were removed in the process of analyzing images. (**C**) Number of cells per image in the 3MM mouse, comparing each location between the base and the tip. There is a significant difference between base and tip for all imaged locations and all 3MM species (T-test, * = p<0.05, ** = p<0.01). Black lines indicate mean. (**D**) Relative bead abundance in different sections of the cecum 220min, 260min, and 300min after gavage of beads, for 5 mice. Beads with different fluorescent labels were gavaged at different time points in the same mouse.

We found that the composition of the microbiota differed in the two locations within the same organ. We found significantly more *B. theta* and *E. coli* at the cecum tip, and more *E. rectale* mostly at the cecum base, with large variation between analyzed images (**Figure 1C**). This indicates that the gut content is not uniformly mixed within a single gnotobiotic cecum. To evaluate whether this intra-cecum heterogeneity can be generalized beyond our gnotobiotic model, cecum tip and base content were harvested from OligoMM^12^ mice containing 12 bacterial species (*34*), and specific pathogen-free (SPF) mice harboring an even more complex microbiota. We determined bacterial community composition in the different parts of the organs using 16S sequencing (**Figure S2A, B**). Differences in community composition were analyzed using Jensen-Shannon distances at the family level (**Figure S2C, D**). While variation in microbiota composition was generally large, family-level Jensen-Shannon distances were significantly larger between parts of the cecum than between cecum tips of the cecum OligoMM^12^ mice (**Figure S2E**), and in SPF mice they were significantly larger between bases than between tips of different ceca (**Figure S2F**). These findings indicate that, in mice with different microbiota complexities, microbiota composition can vary among different parts of the cecum.

Given these differences in bacterial composition at different locations of the cecum, we sought to evaluate how material is transported through the cecum. We orally administered bacteria-sized (1µm diameter) fluorescent beads to SPF mice at three different time points 40min apart. To distinguish beads administered at different times, beads of different colors were administered at every time point. Five hours after the first treatment, the distribution of beads was evaluated (**Figure 1D**). In three out of five mice, the bead abundance in the base dropped monotonously from 220 to 300min, whereas in the other two it increased initially and dropped at the last time point (**Figure 1D**, red bars). The opposite is true for the tip, where relative bead abundance increased monotonously in three out of five animals, while it dropped at 260min and then increasedd at 300min in two (**Figure 1D**, blue bars). Even though bead distribution varied considerably between mice, there are some key insights that can be gained from this data. Our results show a spatiotemporal pattern for particle distribution in the mouse cecum: after entering the cecum lumen at the base, particles moved to the cecum tip on a time scale of hours. The data further suggests that particles move along the epithelium, and only later disperse to the center of the cecum lumen, as relative counts at the tip epithelium increase earlier than counts in the tip center in three out of five mice, and no beads are observed in the tip center in the other two (light vs dark blue bars in **Figure 1D**).

### Bacteria show more microscale clustering than micron-sized beads in the lumen of the murine cecum

We next investigated the spatial structure of the microbiota in the cecum lumen at a micrometer scale. We determined whether bacterial spatial distribution is random, clustered or regular, using the inhomogeneous H(r) function, a derivative of Ripley’s K function (*35*). In short, this method is based on counting the number of cells found in circles of increasing radius r around a focal cell and comparing this to the case in which the distribution is random (i.e., follows a Poisson process; see Methods for details). In our figures, H = 0 indicates random distribution, H > 0 indicates clustering, and H < 0 indicates regularity (**Figure 2A**).

**Figure 2:**
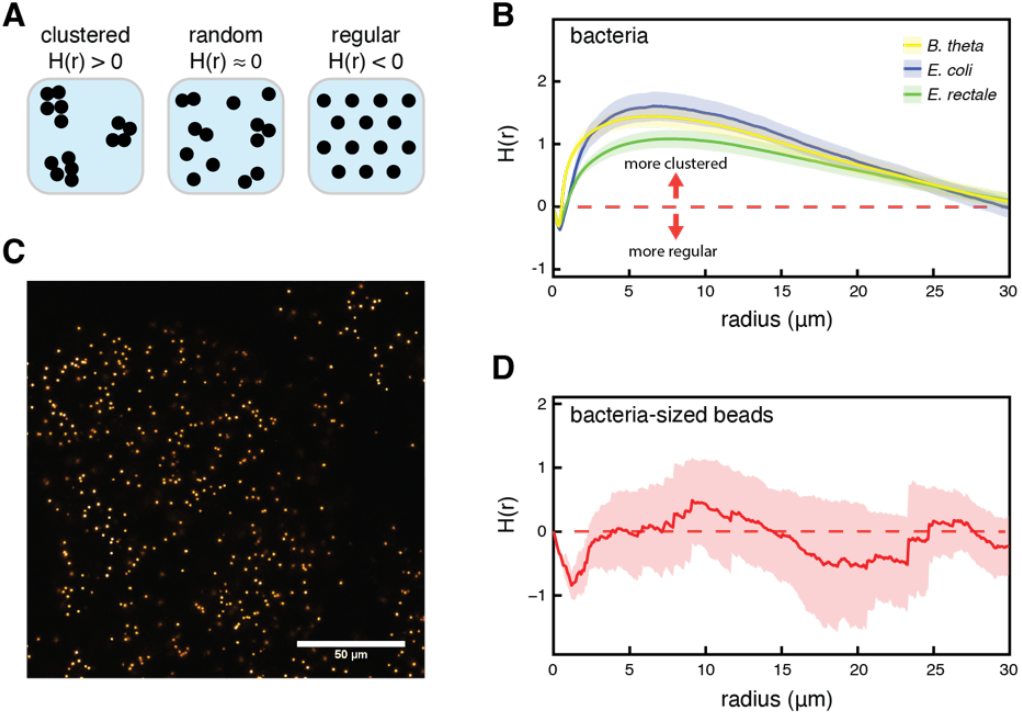
Bacteria display clustering spatial point patterns in the murine cecum. (**A**) Schematic illustration for three possible spatial point patterns: clustered, complete spatial randomness, and regular. (**B**) Results for all three 3MM species when cell distribution was analyzed using the inhomogenous H function. Shaded region indicates 95% CI (n=6). Red dashed line indicates the theoretical value for complete spatial randomness. All bacterial species show significant clustering (p<0.01, Studentized permutation test with Bonferroni correction). Data is shown for all images taken at the cecum base. (**C**) Microscope image of fluorescent beads in the 3MM base center. Beads are 1µm in diameter. (**D**) Results for fluorescent bead distribution in the 3MM mouse cecum, analyzed using the inhomogeneous H function (n=6). Beads are 1µm in diameter, data shown for cecum base. Shaded region indicates 95% CI, dashed red line is the theoretical value for complete spatial randomness. Bacteria-sized beads do not show significant clustering in the 3MM mouse cecum (p=0.736, Studentized permutation test with Bonferroni correction).

We found that all three species of the 3MM were clustered (**Figure 2B**, showing pooled data for all measured locations), regardless of their location in the cecum (**Figure S3**). In addition to a qualitative analysis of clustering, the results of the H function can be employed to estimate the radius of a typical cluster (*35*). Specifically, the value maximizing the H function roughly corresponds to the diameter (or twice the radius) of a typical cluster. Thus, our analysis suggests clusters with radii of around 3-4µm (**Figure 1B**).

Having found that bacteria cluster in the cecum of 3MM mice, we set out to test possible causes of this phenomenon. To test whether abiotic factors, such as micro-scale differences in viscosity or transport dynamics, lead to clustering independent of bacterial activity, we fed 10^9^ 1-µm-diameter fluorescent beads to 3MM mice. These beads approximately mimic the size of single bacteria (**Figure 2C**). We analyzed bead clustering in the cecum 4.5 hours after treatment. In the cecum base, we found no significant clustering of beads (**Figure 2D**; data for cecum base, no beads were found in the cecum tip, likely due to the larger size of the cecum in 3MM mice; when analyzing center and epithelium data separately, beads do exhibit significant clustering at the epithelium, but this clustering is weaker than what we observe for *B. theta* and *E. coli*, **Figure S3B**). This indicates that bacterium-sized, abiotic particles are well mixed on a microscopic scale, at least at the cecum base, and therefore that bacterial activity is necessary for the clustering of cells that we observe in the cecum.

### Antibodies and chemotaxis are not responsible for bacterial cluster formation

As a next step, we tested the role of several biological functions that could play a role in bacterial clustering.

First, we investigated the role of secreted immunoglobulin A antibodies (sIgA) in cluster formation. High avidity sIgA targeting bacterial surface antigens has been shown to enchain bacteria upon division, which can result in bacterial clustering that depends on bacterial growth rates (*22, 36*). We found that ex-germ-free mice colonized for 4 weeks with the 3MM strains produced specific sIgA against *E. coli* and *B. theta*, but not *E. rectale* (**Figure 3A**, red dots). Interestingly, mice which were bred with the 3MM microbiota and were thus colonized from birth had no or very weak specific sIgA production (**Figure 3A**, blue dots), suggesting a role of early life exposure in altering the interaction between gut microbes and the adaptive immune system. Microscopic bacterial counts were similar in 3MM bred mice and ex-germ-free mice, indicating that sIgA did not reduce bacterial population sizes, consistent with the concept that sIgA only drives elimination of bacteria in the context of niche competition (*37, 38*) (**Figure S4A**). When comparing *E. coli* clustering in 3MM-bred mice with ex-germ-free mice colonized for 4 weeks, analysis of the H functions of the images using a Studentized permutation test (*39*) revealed a significant difference between 3MM-bred and ex-germ-free mice (**Figure 3B**), indicating an increase in cluster size and density in the ex-germ-free mice (**Figure S4B**). As we expect bacterial replication and clearance rates in both groups of mice in this experiment to be similar, the observed increase in aggregate size is quantitatively small and likely due to the phenomenon that sIgA-crosslinked clusters are harder to break apart during peristalsis. These dense clusters were visible on microscopy images (**Figure S4B**). However, as clustering is also observed in the absence of detectable IgA responses, we conclude that clustering can occur independently of sIgA.

**Figure 3.**
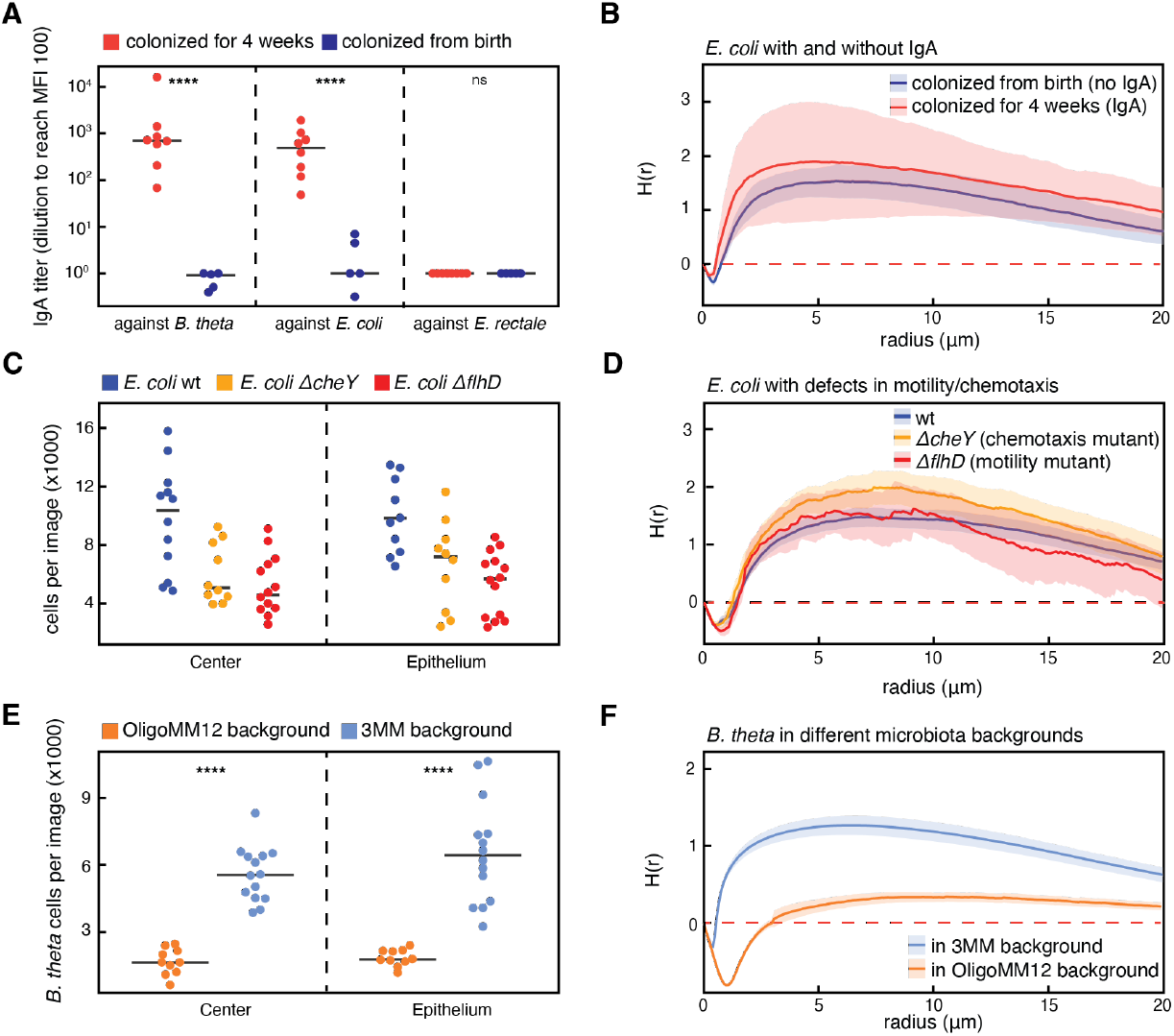
The influence of biotic factors on bacterial cluster formation. (**A**) 3MM-specific IgA titers in mice bred with the 3MM microbiota (n=5) and ex-germ-free mice colonized with the 3MM strains for 4 weeks (n=8). Specific IgA is measured by testing specific binding in small-intestinal lavages against bacterial cultures using flow cytometry. Titers are then calculated by estimating the number of dilutions required to reach a median fluorescence intensity of 100 (T-test, **** = p<0.0001). (**B**) Results for spatial distribution of *E. coli* cells in mice bred with the 3MM microbiota (blue), and ex-germ-free mice colonized with the 3MM microbiota for 4 weeks (red), using the inhomogeneous H function. Both groups show significant clustering (p>0.01), but their respective spatial distributions are not significantly different (p=0.513). Data for cecum tip, center. (**C**) Counts of wild type *E. coli* (wt, blue), chemotaxis mutant (*ΔcheY*, yellow), and motility mutant (*ΔflhD*, red) in ex-germ-free mice on microscopy images. Counts are measured at the cecum base. (**D**) Results for spatial distribution of wt (blue), *ΔcheY* mutant (yellow), and *ΔflhD* mutant *E. coli* cells in the cecum ex-germ-free mice, using the inhomogeneous H function (n=12, 4 in each group, cecum base center data shown). All groups show significant clustering (p<0.01), and there is no significant difference in clustering between groups (p>0.05). (**E**) Counts of *B. theta* cells in the context of the OligoMM^12^ (orange, n=5 mice) and 3MM microbiota (blue, n=6 mice). Data for cecum base (T-test, **** = p<0.0001). (**F**) Results for spatial distribution of *B. theta* cells in OligoMM^12^ (orange) and 3MM (blue) microbiota background, using the inhomogeneous H function. *B. theta* clustering is significantly reduced in OligoMM^12^ as compared to 3MM mice (p<0.01). Shaded regions in **B, D**, and **F** indicate 95% CI, dashed red lines are the theoretical value for complete spatial randomness. Bonferroni-corrected Studentized permutation tests were used for statistical comparisons between H(r) functions.

A second factor that could contribute to bacterial clustering is chemotactic motility (*40*): bacteria might actively congregate following chemotactic cues. To test whether this bacterial trait affect clustering, we colonized germ-free mice with one of three *mCherry* fluorescent *E. coli* strains: one that lacks the ability to perform chemotaxis (Z1331::*cheY*), one that lacks functional flagella (Z1331::*flhD*), and the wild type Z1331 as a control. All three strains colonized germ-free mice (**Figure 3C**), and we evaluated the extent of bacterial clustering ins the cecum in these three cases. Statistical analysis showed no significant difference in clustering between wild type and chemotaxis or flagella deficient *E. coli* cells at the center of the cecum base (**Figure 3D**). At the epithelium of the cecum base, clustering in images with Δ*flhD* mutant bacteria was significantly different than in wild type, suggesting that *E. coli* likely uses flagella in the gut. However, the H(r) function shows stronger clustering in Δ*flhD* than in wild type *E. coli* (**Figure S4C**), indicating that motility is not the origin of clustering, but could rather contribute to cluster dispersal.

### Clustering increases with niche size in the mouse cecum

The next variable that we analyzed as a possible cause of bacterial clustering was the total population size of a given bacterial species in the cecum, the niche size. Given constant loss of bacteria from the GI tract, the relative population size of a given bacterium is a direct readout for the amount of biomass this bacterial population has to produce to maintain a steady state (*41*). While this does not lead to more divisions per cell under steady state conditions, denser populations increase the likelihood of starting clusters closer together. In addition, larger clusters might lead to coordinated loss of the clustered cells, leaving a larger part of the niche open for the remaining cells to fill by growth. Both these phenomena should lead to weaker clustering for bacteria occupying smaller niches. To test this hypothesis, we experimentally decreased the *B. theta* niche relative to 3MM mice by colonizing OligoMM^12^ mice with *B. theta*. Since the OligoMM^12^ microbiota contains 12 species (but not *B. theta*), it presumably contains fewer available niches for *B. theta* to occupy than the 3MM microbiota. Accordingly, *B. theta* counts were >2-fold lower in OligoMM^12^ mice compared to 3MM mice in all studied sections of the cecum, presumably due to a smaller available metabolic niche when competing with the OligoMM^12^ microbiota (**Figure 3E**). While *B. theta* remained clustered, the extent of clustering was significantly reduced in OligoMM^12^ mice as compared to 3MM mice (**Figure 3F**). The results indicate that niche size, and consequently bacterial growth, significantly influence the formation of bacterial clusters in the mouse cecum. This finding suggests that these clusters may represent clonal microcolonies that develop in transiently unmixed cecum content.

### Rheological properties of cecum content could allow microcolony growth between mixing events

To investigate how the physical properties of cecum content and the force exerted by gut contractions might influence microcolony formation, we analyzed the viscoelastic properties of fresh cecum content using rheometry. Our observation that beads traverse the cecum close to the epithelium (**Figure 1D**) could be attributed to a shear-thinning behavior of the cecum content (i.e., decreasing viscosity when force is applied), facilitating movement along the epithelial walls, close to the smooth muscle layer and mucus-secreting goblet cells. To investigate this, we measured the viscoelastic properties of 3MM mouse cecum content. Using a parallel plate, rotational rheometer, we applied oscillatory shear strains at defined amplitudes (the shear strain is defined as the horizontal displacement of the tested material, d, relative to the gap between the plates in the rheometer, g, **Figure 4A**, inset) and a constant oscillation frequency of 1rad/s. The resulting force exerted at every given shear strain (the shear force) is dependent on the material’s viscoelastic properties. We found that shear stress increases with increasing strain amplitude up to a certain threshold, the yield limit. At shear strains higher than the yield limit, shear strain has less effect on shear stress. These findings indicate that cecum content behaves solid-like at low shear stress up to the yield limit (around 60Pa, dashed line in **Figure 4A**), and as a viscoelastic fluid at higher shear stress. In rheology, this behavior is typically illustrated using the storage and loss moduli (G’ and G”, respectively, **Figure 4B**). Conceptually, G’ represents the elastic or stored energy in a material when it is deformed, whereas G” represents the viscous or dissipated energy during deformation. Where these two quantities intersect, the material transitions from more solid-like (i.e., dominated by G’, triangle symbols in **Figure 4B**) to more liquid-like (dominated by G”, circle symbols in **Figure 4B**). As shear forces in cecum content are most likely the result of smooth muscle activity in the cecum wall, this physical property of the cecum content may explain microcolony formation of bacteria: during the time between strong contractions, solid-like behavior creates a matrix that keeps bacteria in place and allows microcolonies to form, while contractions liquify the cecum content and disperse clustered bacteria. This idea is in line with recent findings demonstrating that microbial transport and motility depends on local viscosity (*42*). Additionally, the highest shear strain will likely occur directly at the contact zone between intestinal content and the gut wall, where muscle activity in the gut wall will affect luminal content most strongly. We therefore expect the strongest shear thinning effect in this area, which would lead to low viscosity, consistent with the observed dispersal of beads along the epithelium into the cecum tip (**Figure 1D**).

**Figure 4:**
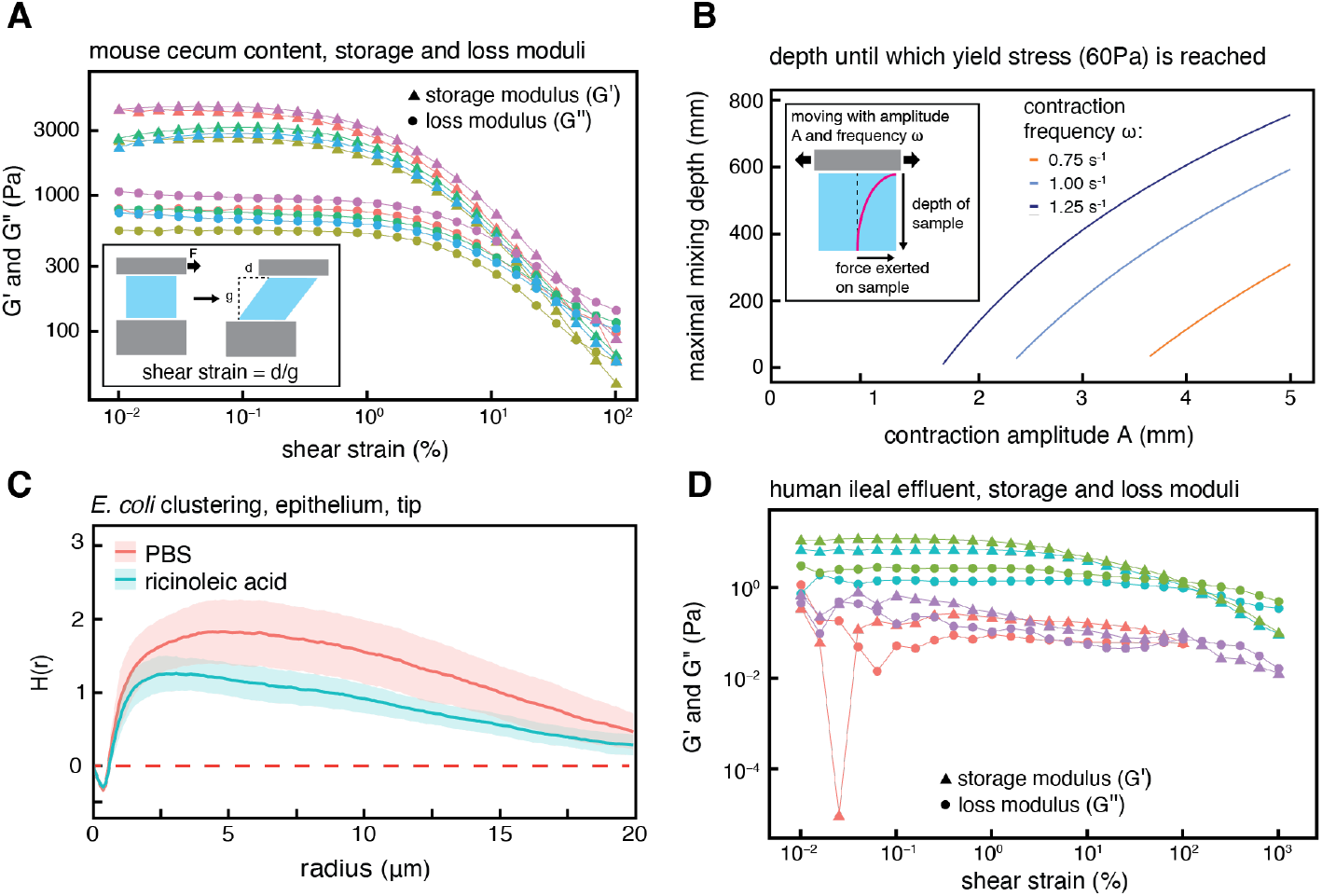
Physical properties of cecum content influence microcolony formation. (**A**) Storage (G’, triangles) and loss (G”, circles) moduli of mouse cecum content. Colors indicate different samples (i.e., cecum content from different mice, n=5). Inset figure illustrates the definition of shear strain, as the fraction of displacement, d, divided by the gap between plates in the rheometer, g. This quantity is dimensionless and usually reported in %. (**B**) Model results based on Stokes’ second problem, predicting the depth a given cecum contraction can reach with enough force to reach the a yield limit of 60Pa (as estimated from our data), depending on the amplitude and frequency (colors) of contractions. Inset image depicts the underlying process. (**C**) Clustering of *E. coli* in 3MM mice treated with ricinoleic acid, a drug increasing smooth muscle activity, is significantly reduced compared to control mice (n=5 per group, p=0.036), while both groups show significant clustering (p<0.01). Shaded areas indicate 95% confidence intervals, red dashed line indicate theoretical values for complete spatial randomness. Data for cecum tip, center. Bonferroni-corrected Studentized permutation tests were used for statistical comparison between H(r) functions. (**E**) Storage (G’, triangles) and loss (G”, circles) moduli of human ileal effluent. Colors indicate different samples (n=4).

Given these rheological properties of cecum content, we wanted to understand whether gut wall movements at realistic amplitudes and frequencies can exceed the yield limit and thus allow dispersion of microcolonies. We combined our bulk rheology data (**Figure 4A, B**) with a simplified model of gut wall movements, based on Stokes Second Problem, to determine the distance from the gut wall at which wall movements are sufficiently strong for shear stress to reach the content’s estimated yield limit of 60Pa, and thus effectively disperse microcolonies. Stokes Second Problem is a classical problem in fluid dynamics that describes the effect of a horizontally oscillating solid surface on an underlying fluid, depending on the fluid’s viscoelastic properties (*43*). Using realistic values for the amplitude and frequency of gut wall movements (see Methods), we estimated the relevant distance from the wall at which the yield stress can be reached to be in the centimeter range (**Figure 4C** and **Figure S5**). While this approach only allows estimating the order of magnitude of the effect of contractions on gut content, it suggests that changing contraction strength could differentially modulate the consistency of content, and thus mixing, throughout the cecum.

To test experimentally whether increased contractile activity in the cecum suppresses microcolony formation, we treated 3MM mice with ricinoleic acid and analyzed bacterial clustering in cecum content 1.5h after treatment. To avoid a direct effect on bacterial activity and growth, the ricinoleic acid was administered by intraperitoneal injection. Ricinoleic acid stimulates smooth muscle activity (*44*), and thus increases contractile activity of the cecum walls. Clustering of *E. coli* and *B. theta* at the epithelium of the tip of the cecum was significantly different and consistently lower in ricinoleic acid treated mice than control mice (**Figure 4D, Figure S6A**), even though *B. theta* tended to be more abundant in mice treated with ricinoleic acid (**Figure S6B**; we found no significant effect of treatment at the center of the cecum tip, and for *E. rectale* at both center and epithelium, **Figure S6C-F**). This data indicates that increased smooth muscle activity of the cecum wall can decrease the extent of bacterial cluster formation.

### Rheological properties of human ileal effluent would allow transient clustering

After finding that the rheological properties of mouse cecum content may facilitate the formation of transient, clonal bacterial clumps between gut contractions, we investigated whether the same phenomenon might take place in the human gut. Analogous to our experiments with mouse cecum content, we measured the viscoelastic properties of ileal effluent of four volunteers with ileostomies and observed qualitatively similar behavior to what we observed in mouse cecum content (**Figure 4E**). We found a diminishing effect of applied shear strain on shear stress, but at values for shear stress that were substantially lower than what we observed in mice, and with more inter-individual variation. G’ is higher than G” at small values of applied shear strain but decreases faster than G” with increasing shear strain (**Figure 4F**), indicating that human ileal effluent, similar to mouse cecum content, transitions from a more solid-like to a more liquid-like behavior when force is applied.

The observed quantitative differences in yield stress between cecum content from mice (around 60Pa, dashed line in **Figure 4A**) and ileal effluent from humans (≈1Pa or lower, slope change in **Figure 4E**) may be attributed to the higher water content of ileal effluent compared to colon content, which has been shown to change when the ileum is disconnected from the colon (*45*). Previous rheology measurements of human feces have found yield stresses spanning a wide range, from 20Pa to 8000Pa (*46*). We expect the rheological properties of human large intestinal content to change considerably along the colon as a function of water resorption, presumably explaining the difference between what we observe for ileal effluent and what has been measured for feces. It is therefore possible that a similar mechanism to the one described in mice could govern microbial assortment in the human large intestine.

## Discussion

Spatial assortment of gut microbiota members is crucial for ecological interactions and evolutionary processes in the gut. In this study, we quantify the spatiotemporal distribution of bacterial cells in the mouse cecum, both on a macroscopic and a microscopic scale, and identify a likely mechanism for the observed behavior.

A first important finding is the distribution of bacteria throughout a whole mouse cecum. We identified a difference in bacterial counts in different parts of the cecum and observed that the main flow from base to tip might be stronger along the epithelium than in the lumen. Subsequent experiments suggest a critical function for the viscoelastic properties of gut content in this phenomenon. Mixing in the mouse cecum generates shear force, which affects gut content most strongly at the epithelium, close to the force’s origin. While there is no structured mucus layer in the mouse cecum, mucus is secreted by the epithelium in the cecum (*47*). Higher mucus concentrations, together with stronger mixing close to the epithelium, could therefore allow for maximum bulk transport of particles in this region, running contrary to the current understanding of mucus as a protective barrier at the epithelium (*48*).

In 3MM mice, and in contrast to SPF mice, small particles did not reach the cecum tip. A possible factor explaining this observation is the larger size of the cecum in 3MM mice, which could result in a more stretched smooth muscle layer that generates weaker contractions (*49*). This could have broad implications for research using gnotobiotic animals, which typically have larger ceca than SPF mice, potentially leading to inhomogeneous distributions of bacterial composition and functions. Future research in the field should therefore consider the heterogeneity in microbial and nutrient composition within the cecum when collecting and analyzing samples, and precise sampling locations should be reported.

Spatial heterogeneity in the gut on a microscopic scale due to biogeographical features, such as proximity to the epithelium or residence in crypts, has been described before (*8–10, 25*), but we found that clustering of bacteria of the same species occurs in the lumen without pre-existing structure. We demonstrated that this can occur independently of antibody-mediated cross-linking and active swimming of cells toward each other and hypothesize that these clusters likely originate from microbial growth in conjunction with the rheological properties of cecum content: without active mixing, cecum content behaves as a thick gel, holding bacterial offspring in place after cell division. A similar mechanism for microcolony formation has been shown for S. aureus entrapped in agarose gels (*50*), and research in zebrafish has described clustering of gut bacterial and suggested that gut contractions might play a role in limiting cluster size (*25*). We have now demonstrated this in mice and found evidence that human gut content has rheological properties that could allow the same phenomenon in the human colon. Formation of bacterial clusters in the intestinal tracts of animals that is driven by growth and opposed by gut contractions might therefore be a general phenomenon.

Bacterial clustering at the micro-scale can have functional consequences. Metabolic interactions between bacteria happen at very short ranges (*12, 51*), and spatial arrangement thus plays a crucial role for allowing or blocking ecological crosstalk between bacterial cells. In addition, the duration of spatial assortment can affect whether interactions between cells can play a role in evolutionary processes. Gut contractions which disperse clonal microcolonies can therefore play an important role in maintaining or disrupting interspecies interactions, giving the host a direct way to shape ecological and evolutionary processes in the gut microbiota. Micrometer-scale clustering, similar to what we observe in the mouse cecum, has been shown to lead to changes in antibiotic susceptibility in nutrient replete conditions in vitro (*50*), and gel-entrapped microbes generally are known do exhibit similarly reduced susceptibiltiy to treatment with antimicrobials as bacterial cells growing in biofilms (*52–55*).

Our study highlights the importance of quantitatively understanding the spatiotemporal dynamics of bacterial activity in the gut microbiota. Using image analysis in combination with statistics on spatial point patterns, we found that bacteria form microcolonies in the gut. By investigating how microbial behavior is affected by the physical properties of its environment, we were able to determine the relative importance of various factors that might contribute to this phenomenon. We conclude that bacterial microcolony formation is largely influenced by two competing properties, bacterial growth and gut contractions, which likely play an important role in gut microbial ecology.

### Lead contact

Further information and requests for resources and reagents should be directed to and will be fulfilled by the lead contacts, Markus Arnoldini or Giorgia Greter.

## Materials and Methods

### Experimental design

The research objective of this study was to determine the causes of local spatial structure in the murine cecum and estimate the extent of microbial spatial structure in the human large intestine. The overall design used controlled laboratory experiments and utilized a few human samples to estimate parameters of physical properties of the human intestine.

Wild type C57BL/6 mice were maintained on a standard chow diet at the ETH Phenomics center. Mice were re-derived Germ-free or bred with the 3MM or OligoMM^12^ microbiotas and kept in isolators. The 3MM microbiota was derived using strains *Bacteroides thetaiotaomicron Δtdk* (*56*) (a strain that behaves like wild type, but allows counterselection for genetic engineering, derived from ATCC29148), *Eubacterium rectale* ATCC33656, and *Escherichia coli* HS (*57*). Specific pathogen free (SPF) mice were bred and housed in individually ventilated cages. All experiments began at 8-14 weeks of age. Mice were fed *ad libitum* for the duration of all experiments and were kept with a dark and light period of 12 hours each. All animal experiments were approved by the Swiss Kantonal authorities (Licenses ZH120/19 and ZH016/21).

For each experiment, a minimum sample size of n=4 per experimental group was selected. For experiments in which mice were colonized with bacteria, an endpoint was selected of 2 days post colonization, allowing bacteria to reach a steady state colonization levels in the cecum.

### DNA isolation and 16S sequencing

Cecum content was collected from 3MM, OligoMM^12^, and SPF mice at the tip and the base. Bacterial DNA was extracted using the DNeasy PowerSoil Pro Kit 420 (Qiagen, Germany). 16S DNA sequencing was performed with Biomarkers Technologies (BMKGENE). Illumina sequencing was performed using an Illumina NovaSeq 6000, with paired end reads and read length of 100,000 (Illumina).

The raw sequencing data were analysed using DADA2 v1.14 (*58*). Specifically, primer sequences were removed with cutadapt v2.8 (515F = GTGCCAGCMGCCGCGGTAA, 806R = GGACTACHVHHHTWTCTAAT) and only inserts that contained both primers and were at least 75 bases long were kept for downstream analysis (*59*). Next, reads were quality filtered using the filterAndTrim function of the dada2 package (maxEE=2, truncQ=3, trimRight = (40, 40)). The learnErrors and dada functions were used to calculate sample inference using pool=pseudo as parameter. Reads were merged using the mergePairs function and bimeras were removed with the removeBimeraDenovo (method=pooled). Remaining amplicon sequence variants were then taxonomically annotated using the IDTAXA classifier in combination with the Silva v138 database (*60, 61*).

### Calculating Jensen-Shannon divergence between microbiomes

For each sample, the relative species abundance based on 16S sequencing data can be interpreted as a probability distribution on the set of bacteria. The Jensen-Shannon divergence is a way to compare two probability distributions, with low/high values indicating similarity/dissimilarity. If *B* is the set of bacterial strains, and *P* and *Q* are two distributions on *B*, the Jensen-Shannon divergence between *P* and *Q* is defined as

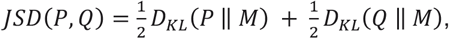

where 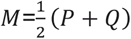 is a mixture distribution of *P* and *Q*, and *D*_*KL*_ is the Kullback-Leibler divergence defined as

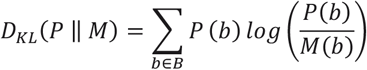

While the Kullback-Leibler divergence is asymmetric, the Jensen-Shannon divergence is symmetric. We computed the Jensen-Shannon divergence for each sample pair, with results shown in **Figure S2C** and **D**. Samples were categorized into base-samples and tip-samples. We then aggregated the Jensen-Shannon divergence values from each base sample to all other base samples (forming the base/base group) and from each tip sample to all other tip samples (forming the tip/tip group). Additionally, we calculated the divergence from base samples to tip samples, creating the base/tip group. A one-sided Mann-Whitney U test was conducted to assess differences in distances between these groups, as illustrated in **Figure S2E** and **F**. For instance, in SPF mice, base samples exhibited greater similarity to one another compared to the tip samples, a trend not observed in OligoMM^12^ mice.

### Fluorescent in situ hybridization

Following euthanasia, the entire mouse cecum was harvested for microscopy. Freshly harvested samples were submerged in 4% paraformaldehyde and incubated overnight at 4°C. Samples were then transferred to a 20% sucrose solution and incubated for 4 hours at room temperature. After fixation, the samples were cut into tip and base sections and embedded in O.C.T (Tissue-Tek®, O.C.T.™ Compound), snap frozen in liquid nitrogen and stored at -80°C. Cecum tip and base were cut into 5 μM thick sections using a Microm HM525 cryotome (ThermoFischer Scientific). The sections were mounted on a Superfrost++ glass slides and stored at – 20°C for further use.

Fluorescent In Situ Hybridisation (FISH) was used to stain each 3MM member. Dried tissue sections were contoured with a hydrophobic pen (ImmEdgeTM, Vector laboratories). Subsequently, samples were washed with PBS, dehydrated with increasing concentrations of ethanol (50%, 80%, 100%, vol/vol) and air dried. Five probes (one for *E. rectale*, one for *E. coli*, and three for *B. theta*) were used to detect the 3 bacterial species and were diluted in the hybridization buffer (for 100ml: 10ml 5M NaCl, 2ml 1M Tris-HCl, 0.1ml 10% SDS, 10ml formamide, 69.9ml H2O) to a final concentration of 1% (vol/vol) each. The probe mix was added to the sections and incubated for 4 hours in a dark moist chamber at 50°C. Washing buffer (for 100ml: 10mL 5M NaCl, 2ml 1M Tris-HCl, 0.1ml 10% SDS, 79.9ml H_2_O) was added to the sections and incubated for 20 minutes at 50°C. Three additional washing steps of 10 minutes were performed with PBS. After washing, the slides were rinsed with ice cold ddH2O, air dried and mounted with Vectashield® H-1400TM Hardset Antifade mounting medium (Vector Laboratories). Slides were stored at -20°C for further use.

### Niche size experiment

To evaluate the effect of niche size on bacterial clustering, an mCherry fluorescent *B. theta* strain was used (*62*) to colonize OligoMM^12^ mice. Mice were colonized for 3 days and subsequently euthanized. Entire cecum was collected, fixed, cut into sections, and mounted as described above.

### Motility experiment

To evaluate the effect of bacterial motility on bacterial clustering, *E. coli* Z1331 wild-type and mutant strains were used (*63, 64*). Gene deletions (*cheY*::*aphT* or *flhD*::*aphT*) in *E. coli* were obtained via PCR-based inactivation of chromosomal genes, as previously described (*65*). Ampicillin resistance and *mCherry* fluorescence genes were inserted using plasmids pSIM5 and pFPV25.5 mCherry (*47*). Germ-free mice were colonized with *E. coli, E. coli cheY*::*aphT*, or *E. coli flhD*::*aphT*. Mice were colonized for 3 days and subsequently euthanized. Entire cecum was collected, fixed, cut into sections, and mounted as described.

### Particle mixing experiment

To evaluate whether small particles are mixed in the mouse cecum, 6 3MM mice were treated by oral gavage with 100µL of 10^9^ Fluoresbrite® Yellow Green 1.0µM Microspheres (Polysciences). Mice were euthanized 4.5 hours after treatment and entire cecum was collected, fixed, cut into sections, and mounted as described.

To determine the flow of small particles in the mouse cecum, SPF mice were treated by oral gavage three times, 40 minutes apart with 50µL of 10^9^ Microspheres. The first treatment was with Fluoresbrite® Polychromatic red 1.0µM Microspheres (Polysciences), followed by Fluoresbrite® Polyfluor® 407 1.0µM Microspheres (Polysciences), and finally with Fluoresbrite® Yellow Green 1.0µM Microspheres (Polysciences). Mice were euthanized 5 hours after the first treatment with polychromatic red microspheres and entire cecum was collected, fixed, cut into sections, and mounted as described.

### Manipulating gut contractions

To evaluate the effect of gut contractions on bacterial clustering, 3MM mice were treated with ricinoleic acid or PBS to increase gut contractions or as a negative control, respectively. 8mg of ricinoleic acid were dissolved in 100µL of PBS and sterile filtered. Mice were injected intraperitoneally with 100 µL of PBS or diluted ricinoleic acid, and euthanized after 90 minutes, based on previous work (*44*). The cecum was fixed, sectioned, stained using FISH, and mounted as described.

### Image acquisition and processing

Acquisition of bacterial images in the cecum was performed using a Leica TCS SP8 STED confocal microscope under a HC PL APO CS2 63x/1.4 oil immersion objective. The *E. coli* probe, tagged with an Atto 425 fluorophore, was excited with a 458 nm argon laser and emission signals were detected between 471/539 nm. A DPSS 640 nm laser was used to excite the *B. theta* probes and a DPSS 471 nm laser to excite the *E. rectale* probe. The emission was detected in a 650/712 nm and 576/618 nm window, respectively. Sections of cecum containing *mCherry* fluorescent bacteria were excited with a 458 nm argon laser and emission signals were detected between 471/539 nm.

For fluorescent beads in particle mixing experiments, Fluoresbrite® Polyfluor® 407 1.0µM Microspheres were excited with a 405 nm argon laser and emission signals were detected between 415/440 nm. Fluoresbrite® Yellow Green 1.0µM Microspheres were excited with a 458 nm argon laser and emission signals were detected between 468/500 nm. Fluoresbrite® Polychromatic red 1.0µM Microspheres were excited with a 514 nm argon laser and emission signals were detected between 524/600 nm.

Images were acquired with a resolution of 2048×2048 pixels. Ten Images were acquired as Z-stacks across the 5 μM thick section. After acquisition, images were deconvoluted using the Huygens Remote Manager. A signal to noise ratio of 9 was used for the deconvolution of *E. coli*, and signal to noise ratio of 17 was used for the deconvolution *E. rectale* and *B. theta*. The deconvolution algorithm “cmle” was used in 30 iterations. The resulting deconvoluted stacks were compressed into one layer using a maximum intensity projection in ImageJ (v 1.51n) and each channel image was stored in a separate folder.

Four images of each mouse cecum section were taken, two at the epithelial border and two in the center of the cecum. The images covered a total area of 184.5 μm^2^. For the experiment evaluating fluorescent bead mixing in 3MM mice, the resolution was decreased to 1024×1024 pixels to increase image acquisition speed.

For the experiments evaluating *E. coli* mixing through IgA-mediated clustering in 4-week colonized mice, as well as for evaluating small particle flow direction in the cecum in SPF mice, two images of each mouse cecum section were taken: one at the epithelial border and one in the center of the cecum. Epithelial images were composed of 6 square images stitched into a line and center images were acquired as 9 single images stitched into a larger square. Epithelium images covered a total area of 0 ∼85×510 μm and center images an area of ∼255 μm^2^.

### Image analysis

An image analysis pipeline was developed in Matlab (R2022a) to identify the coordinates of each bacterium in microscopy images. Briefly, image preprocessing was done using top hat filtering and local maxima adjustments. A binarization threshold was set based on the average pixel intensity in each image and single cells were identified using size, eccentricity, and solidity thresholds. Food particles were removed from the images based on size and length/width ratios. Cells in clusters were identified two times by edge detection and subsequent dilation/erosion. The total cell count per image, coordinates, and area of each bacterium in the image was stored for further analysis.

### H function

The inhomogenous H function was used to evaluate the spatial point patterns in all images. Ripley’s H function is a transformation of Ripley’s inhomogenous K function, in which the number of neighbors of bacteria (*K*_*inhom*_) are calculated at different radii (r) from all cells and compared to simulations of complete spatial randomness. The inhomogeneity of bacterial density in images due to food particles inhibiting the occurrence of bacteria is accounted for by weighing each point to its local density. The formula used in Ripley’s inhomogenous K-function is as follows:

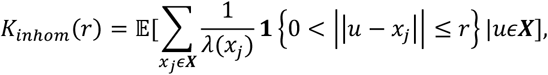

defined as the expected number of neighboring points *x*_*j*_ within a radius *r* of a focal point *u* of the point process ***X***, weighted by the local density at that point *λ*(*x*_*j*_).

The L-function transforms the K-function such that images with point patterns of complete spatial randomness will have *L*_*inhom*,_(*r*) = *r* for all distance bands, while images with clustered bacteria will have *L*_*inhom*,_(*r*) > *r*, and images with regularly spaced bacteria will have *L*_*inhom*,_(*r*) < *r*. The L-function is as follows:

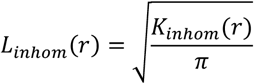

The H function is a normalization of *L*_*inhom*,_(*r*), such that all values of complete spatial randomness would be *H*_*inhom*,_(*r*) = *L*_*inhom*,_(*r*) − *r* = 0. All images with clustered bacteria would be *H*_*inhom*,_(*r*) > 0, and all images with regularly distributed bacteria would be *H*_*inhom*,_(*r*) < 0. Using this transformation, we could visualize at which radius clustering is occurring based on the curve’s deviations from 0.

To test whether the *H*_*inhom*,_(*r*) curves were significantly different from each other, a Studentized permutation test was carried out with 1000 permutations over an analysis radius of 0 to 20 µm (*39*). For calculating H(r) functions, plotting, and associated statistical analysis, the R-package “spatstat” V4.3.2 (*66*) was used. Specifically, we employed the *Linhom* function with the parameters *correction=c(“border”), renormalise=FALSE, ratio=TRUE*, while maintaining the standard settings for all other options.

### IgA measurements

Wild-type GF mice were colonized with the 3MM microbiota for 4 weeks. Intestinal lavage was collected in 2mL of PBS from ex-GF mice as well as from WT mice bred with the 3MM microbiota and stored at -80°C for further analysis. A flow cytometry (FACS) technique was applied to estimate 3MM specific IgA responses (*67*). Intestinal lavages were diluted 5 times in 3 folds across a 96-well plate, starting from undiluted samples. The 3MM bacteria were grown overnight in filtered BHIS, washed, and quantified by flow cytometry. Subsequently they were diluted in PBS containing 2% BSA and 0.02% NaN3 to a concentration of roughly 10^5^ bacteria and added to each intestinal lavage dilution. BrilliantViolet 421 anti-rat IgA antibody (BD biosciences) was used to detect IgA. Plates were read by a CytoFlex Flow Cytometer (Beckman Coulter). Bacteria were identified by light scattering methods and IgA bound to bacteria by median fluorescent intensity (MFI) using FlowJo™ V10.8 Software (BD Life Sciences).

### Rheology of human samples

Stoma effluent samples were collected from 3 ileostomy patients of the NBMISI (Networks of bacterium-metabolite interactions in the small intestine) study. Samples were taken by the study physician with a sterile syringe directly from the stoma bag. Samples at timepoint 0h are collected after an overnight fasting period. Subsequent timepoints indicate collection times after a defined nutritional intervention (Nutricia Calogen® or Nutricia preOp®, respectively after an overnight fasting period); e.g., timepoint 3h indicates sampling at 3 hours after ingestion of a nutritional intervention. No further food was ingested after the nutritional intervention. Water intake was assessed but not restricted after the nutritional interventions.

The NBMISI study (ClinicalTrials.gov ID: NCT04978077) operates with the approval of the responsible local ethics committee (KEK Bern, project ID 2021-01108, approved on October 20th, 2021). Only patients who provided written informed consent were included in the NBMISI study.

### Rheology

To measure the viscoelastic properties of mouse cecum content, the entire cecum content was harvested from 5 3MM mice and stored at -80°C for further analysis. Human ileostomy samples were also stored at -80°C until rheology measurements were performed.

The viscoelastic response of each sample was quantified using an Anton Paar MCR302 rheometer. To minimise slip during the rheology measurement, grit paper was added to the bottom plate and the measurement geometry of the rheometer using double sided tape. All measurements were performed using a parallel plate geometry with 25 mm diameter. For each measurement, the temperature was set at 37°C, and humidity was maintained by adding a wet sponge to the rheometer closure. Oscillatory amplitude sweeps were performed at a frequency of 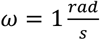 and strain range of *γ* = 0.1 *to* 1000%. Gap size was 0.8mm. Frequency sweeps were performed at strain of *γ* = 0.1% and a frequency range of *ω* = 0.1 *to* 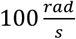.

### Contraction depth modelling

We modelled the gut wall contractions and the corresponding mixing process as the ‘Stokes second problem’, a classic fluid dynamics problem, whereby the wall contractions are represented as an oscillating plane (in-plane oscillations) that mixes the fluid (digesta) above it. We approximated the digesta as a Newtonian fluid of density *ρ* and dynamic viscosity *μ*. Digesta density *ρ* was determined by measuring weight and volume changes simultaneously when digesta was added to water (results: 1.16g/ml, 1.29g/ml, 1.31g/ml, for three independent replicates). The rigid plane is located at *y* = 0 and oscillates with velocity *Aω* cos (*ωt*) along the *x* -direction, where *A* and *ω* are the amplitude and frequency of the oscillations. The digesta velocity profile then reads

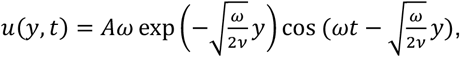

where *ν* = *μ*/*ρ* is the kinematic viscosity of the digesta. The shear stress is

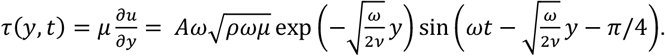

We first compute the shear stress at *y* = 0, which represents the force per unit area the gut wall exerts on the digesta

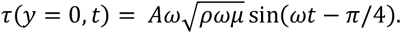

The amplitude of the force is thus 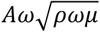, which is plotted in **Figure S5** as a function of the amplitude *A* for various oscillation frequencies *ω*.

To estimate the mixing depth *D*, we match the amplitude of the viscous shear stress at *y* = *D*, with the yield stress *τ*_*yield*_

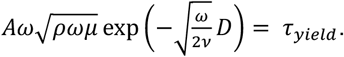

Solving for *D*, gives

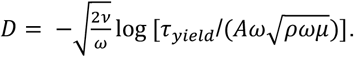

For producing the plots in **Figure 4C**, we estimated the following parameters to solve for D:

| Parameter | Value | Source |
| --- | --- | --- |
| Digesta yield stress $\tau_{yield}$ | 60 Pa | rheology, this study |
| Digesta dynamic viscosity $\mu$ | 2237 Pa s | rheology, this study |
| Digesta density $\rho$ | 1250 kg/m<sup>3</sup> | density measurements,<br>this study |

## Acknowledgements

We would like to acknowledge Jan Vermant and Jonas Cremer for critical reading of the manuscript, and all members of the laboratory for Mucosal Immunology at ETH Zürich for their input. We thank Sven Nowok and Dominik Bacovcin, as well as all the animal caretakers at the EPIC facility at ETH Zürich for maintaining the mouse lines and for their experimental support.

## Author Contributions

Conceptualization: GG, ESl, MA

Experiments and data analysis: GG, SH, PB, DK, ND, NR, SGa, CM, SGe, LL, WDH, MA

Human sample collection: SJ, BM, BY

Mathematical models: MR, ESe, JS, RS

Writing: GG, MA

Supervision: MA, ESl

## Data and materials availability

All data relevant for generating figures is available in a curated data archive at ETH Zurich under the DOI (https://doi.org/10.3929/ethz-b-000665345). All code used for data analysis and for generating figures is available on Gitlab under the link (https://gitlab.ethz.ch/ggre/mixing-project-scripts). All data was analyzed and plotted using R V4.3.2.

## Supplementary figures

**Figure S1.**
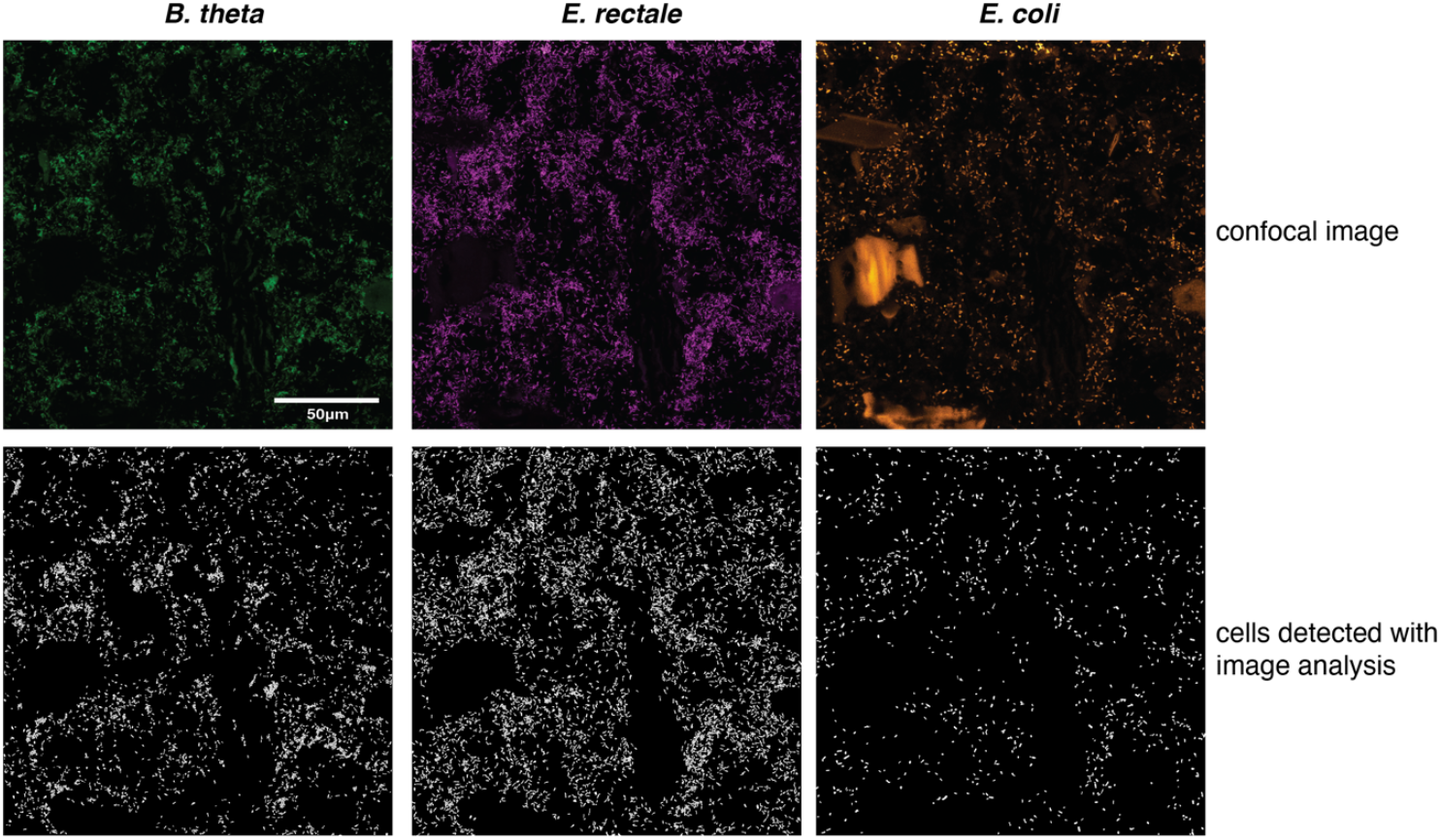
Example for cell detection using the image analysis pipeline developed for this study. Top row shows confocal microscopy images for the three bacterial species in the 3MM microbiota, detected by FISH probes in fixed cecum content. Bottom row shows the binary images resulting from image analysis.

**Figure S2.**
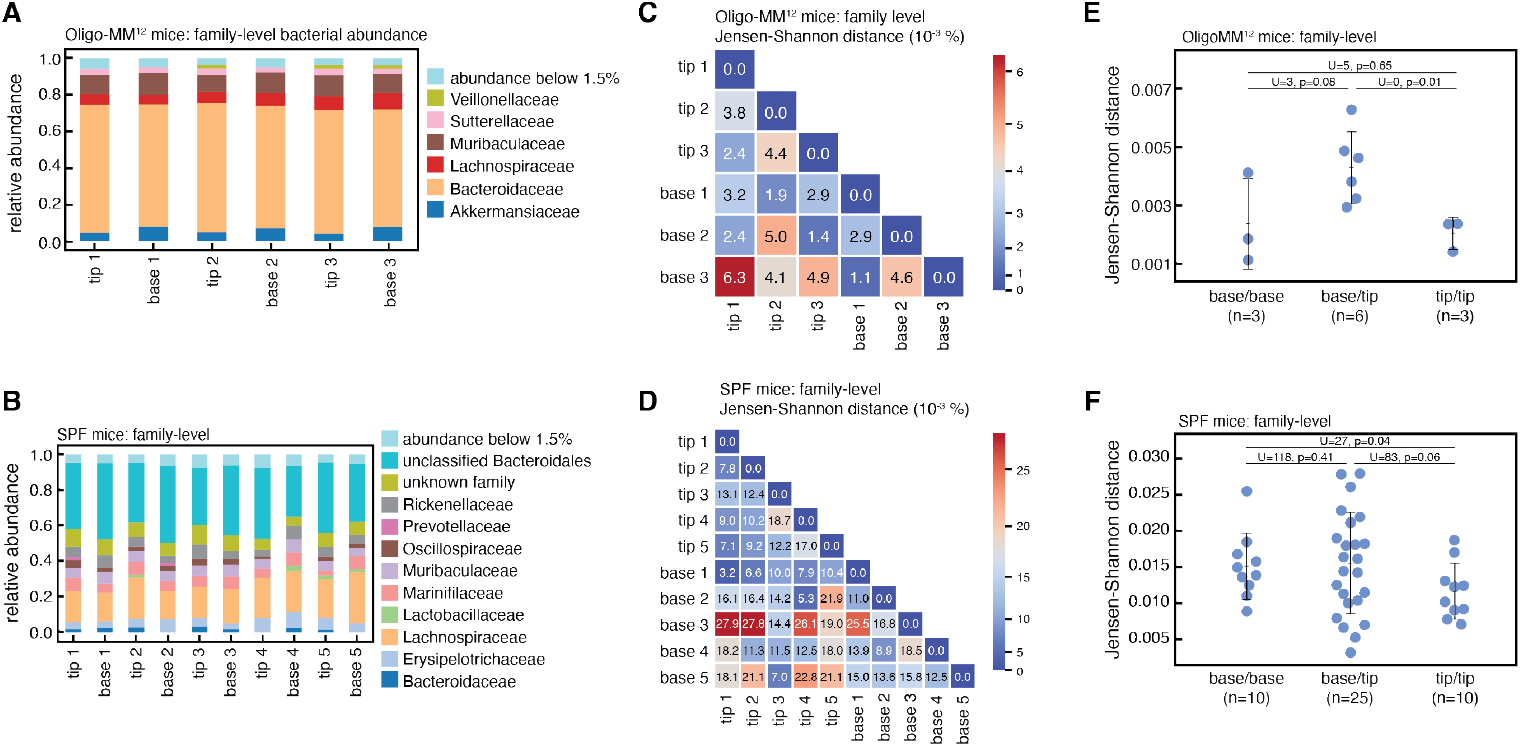
The microbiota is not well-mixed in the cecum of SPF and OligoMM^12^ mice. (**A**) Results of 16S sequencing of cecum content of OligoMM^12^ (n=3) and (**B**) SPF mice (n=5) at the family level. Tip and base samples of the same cecum were sequenced separately. (**C**) Jensen-Shannon distances between each sequenced tip and base for OligoMM^12^ and (**D**) SPF mice at the family level. (**E**) Plots of base/base, base/tip, and tip/tip Jensen-Shannon distances across all mice in the OligoMM^12^ group and (**F**) the SPF group. Significance was tested with a one-sided Mann-Whitney U test.

**Figure S3.**
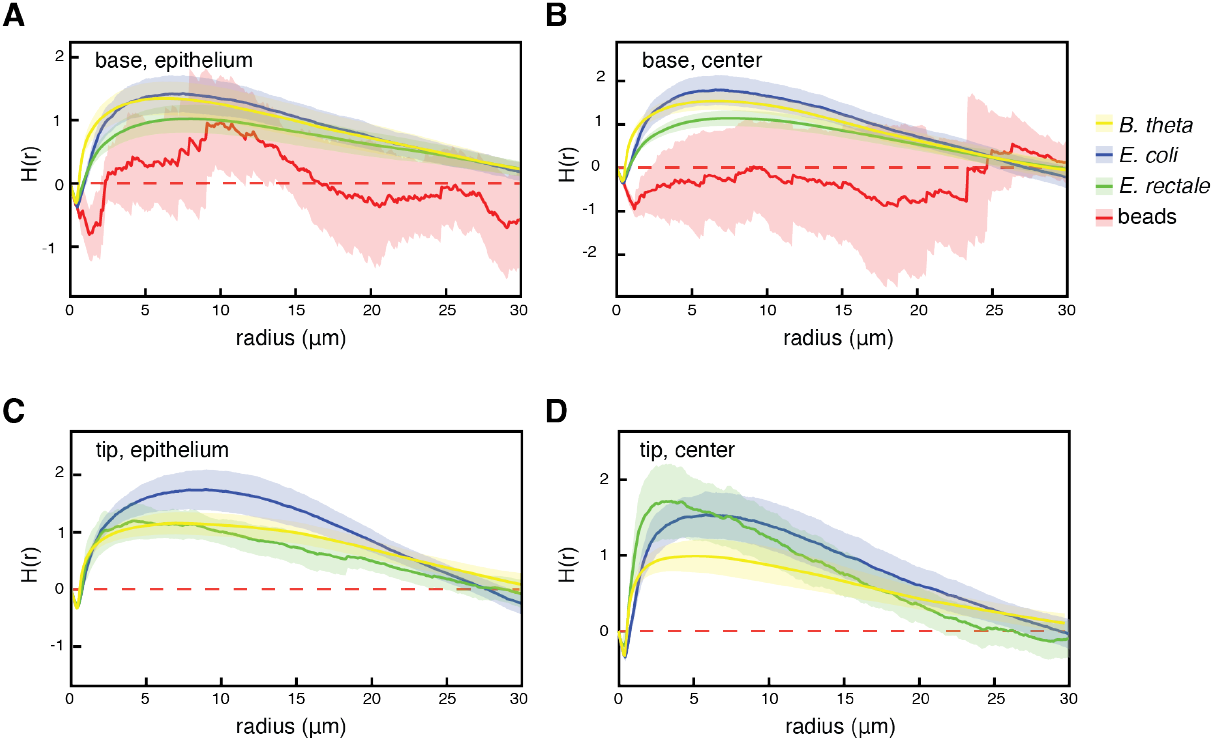
3MM microbiota members, but not beads, cluster in the cecum lumen. Results for all three 3MM species when cell distribution was analyzed using the inhomogeneous H function. Shaded regions indicate 95% CI (n=6). Red dashed lines indicate the theoretical value for complete spatial randomness. Locations are (**A**) base, epithelium, (**B**) base, center, (**C**) tip, epithelium, and (**D**) tip, center. H(r) functions for all bacterial species are significantly different from complete spatial randomness (p<0.01). With the exception of *E. rectale* at the epithelium of the cecum base, H(r) functions for all bacteria data were significantly different from H(r) function for beads (p<0.01). H(r) functions for beads are significantly different from complete spatial randomness at the base epithelium (p=0.04), but not at the base center. No beads were found at the cecum tip. Bonferroni-corrected Studentized permutation tests were used for statistical comparisons between H(r) functions.

**Figure S4.**
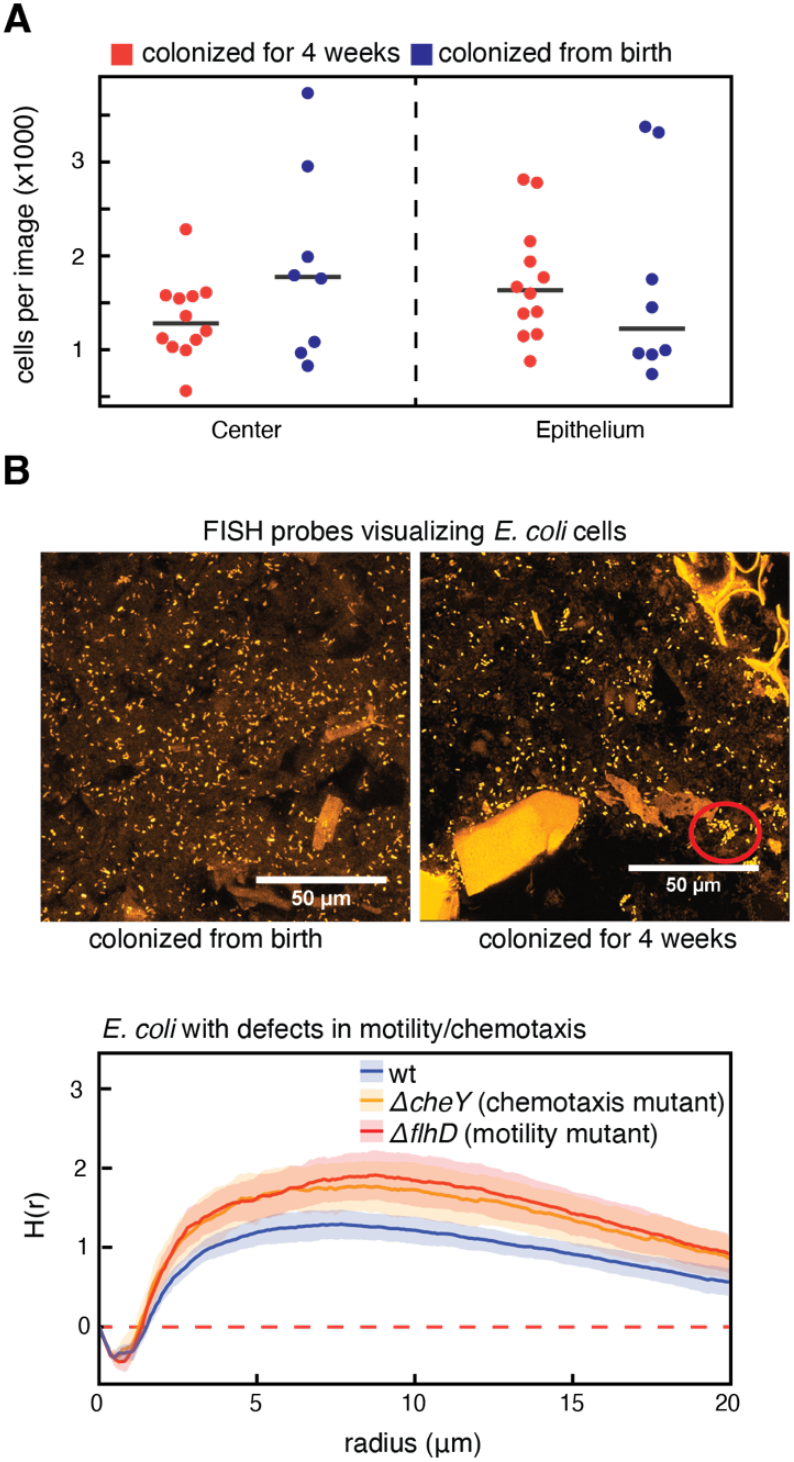
Effects of sIgA on bacterial colonization and visual cluster formation. (**A**) Cell counts, based on image analysis, for E. coli in mice that were colonized from birth with the 3MM microbiota (blue), and mice that were colonized for 4 weeks (red). Only mice that were colonized for 4 weeks have sIgA antibodies against *E. coli*. (**B**) Representative microscopy images with FISH-stained *E. coli* cells in fixed cecum content, from mice colonized from birth (left) and mice colonized for 4 weeks (right). A tight cluster that is presumably formed by sIgA-cross-linking is marked by a red circle in the image on the right. (**C**) Results for spatial distribution of wt (blue), *ΔcheY* mutant (yellow), and *ΔflhD* mutant *E. coli* cells in the cecum ex-germ-free mice, using the inhomogeneous H function (n=12, 4 in each group, cecum base epithelium data shown). All groups show significant clustering (p<0.01), and there is no significant difference in clustering between wild type and Δ*cheY* (p>0.05), but between wild type and Δ*flhD* (p=0.12). The form of the H(r) function suggests that Δ*flhD* clusters more than wild type, indicating that flagella might be important for cluster dispersal rather than cluster formation. Bonferroni-corrected Studentized permutation tests were used for statistical comparison between H(r) functions.

**Figure S5.**
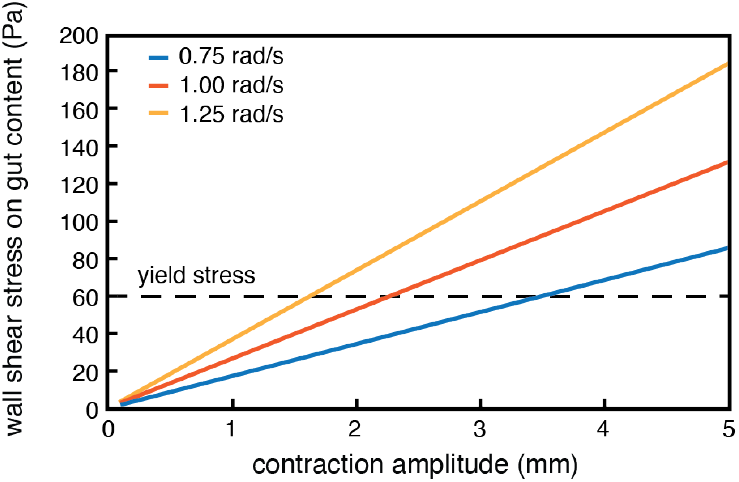
Dependence of shear stress experienced by gut content on the amplitude and frequency of gut contractions. The shown results are based on Stokes’ second problem and take measured values for viscosity and density of gut content into account. Dashed line indicates measured yield stress of gut content.

**Figure S6.**
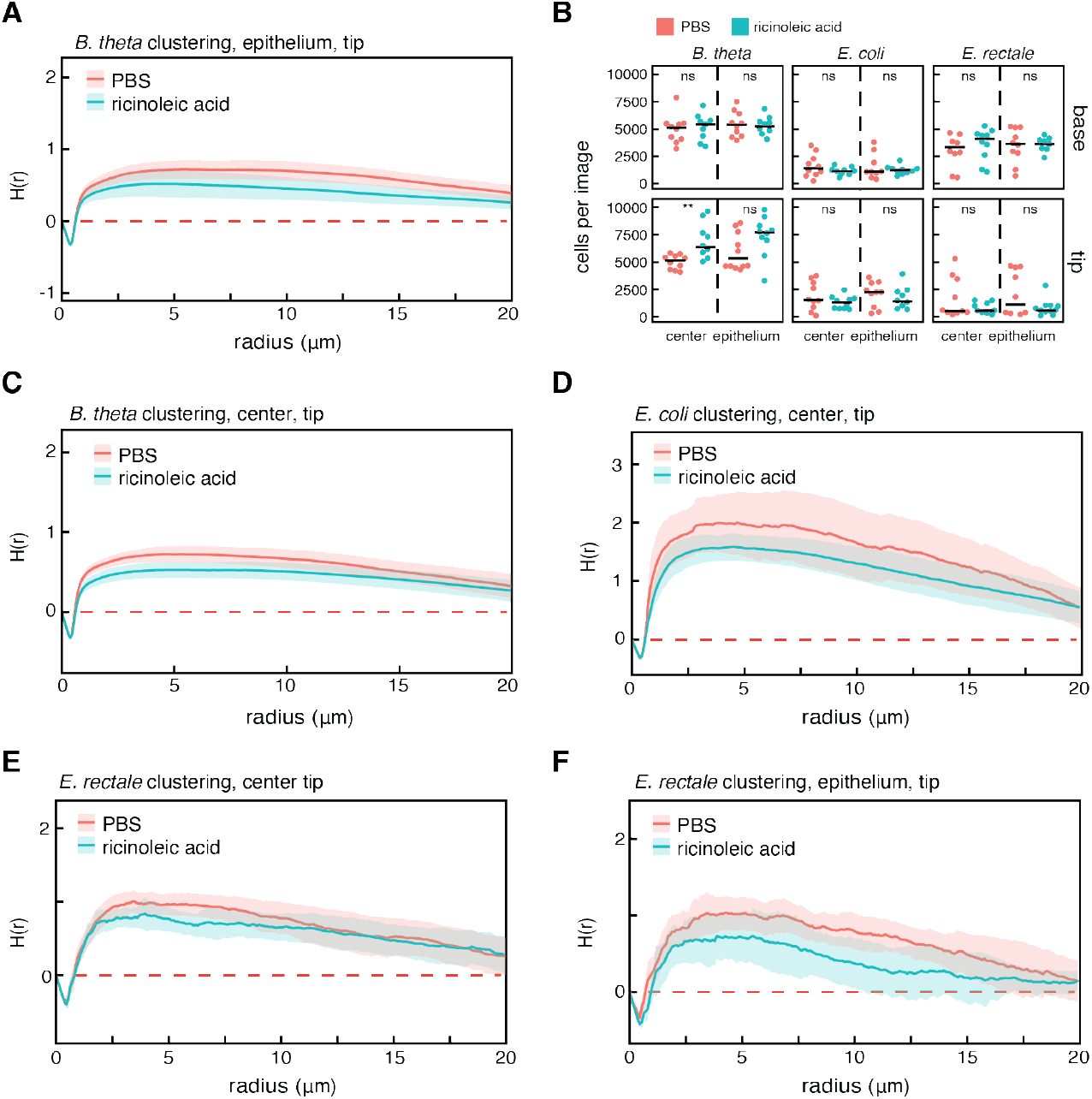
Analysis of clustering and population sizes under modified gut contractions. (**A, C-F**) Clustering analysis using the inhomogeneous H function for all three 3MM strains after IP injection of ricinoleic acid (green) or PBS (red), for different locations in the cecum. Studentized permutation test shows that the datasets in (**A**) are significantly different, whereas (**C-F**) are not. However, in all ricinoleic acid treated groups a clear trend towards lower clustering is visible. (**B**) Cell counts on microscopy images in ricinoleic acid and PBS treated groups. B. theta is significantly less abundant in the center of the cecum tip in PBS treated vs ricinoleic acid treated mice. All other cell counts are not significantly different (T-test, ** = p<0.01).

